# Hookworm-derived small molecule extracts suppress pathology in a mouse model of colitis and inhibit secretion of key inflammatory cytokines in primary human leukocytes

**DOI:** 10.1101/316885

**Authors:** Phurpa Wangchuk, Constantin Constantinoiu, Konstantinos A. Kouremenos, Luke Becker, Linda Jones, Catherine Shepherd, Geraldine Buitrago, Paul Giacomin, Norelle Daly, Malcolm J. McConville, Rachael Y. M. Ryan, John J. Miles, Alex Loukas

**Author notes:** Corresponding author: Prof. Alex Loukas, Australian Institute of Tropical Health and Medicine, Building E4, James Cook University, McGregor Rd., Smithfield, QLD 4878, Australia; Dr Phurpa Wangchuk, Australian Institute of Tropical Health and Medicine, Building E4, James Cook University, McGregor Rd., Smithfield, QLD 4878, Australia.

## Abstract

Iatrogenic hookworm therapy shows promise for treating disorders that result from a dysregulated immune system, including inflammatory bowel disease (IBD). Here we use a metabolomics approach to characterize the non-protein small molecule complement of hookworms. Gas chromatography-mass spectrometry and liquid chromatography-mass spectrometry analyses of somatic tissue extracts revealed the presence of 52 polar metabolites and 22 non-polar components including short chain fatty acids (SCFA). Several of these small metabolites, notably the SCFA, have been shown to have anti-inflammatory properties in various diseases, including IBD. Using a murine model of colitis and human peripheral blood mononuclear cells, we demonstrate that somatic tissue extracts of the hookworm *Ancylostoma caninum* contain small molecules with anti-inflammatory activities. Of the five extracts tested, two of them significantly protected mice against T cell-mediated immunopathology and weight loss in a chemically-induced colitis model. Moreover, one of the anti-colitic extracts suppressed *ex vivo* production of inflammatory cytokines from primary human leukocytes. While the origin of the SCFA (parasite or host microbiota-derived) present in the hookworm somatic tissue extracts cannot be ascertained from this study, it is possible that *A. caninum* may be actively promoting an anti-inflammatory host microbiome by facilitating immune crosstalk through SCFA production.

---

Inflammatory Bowel Disease (IBD) is associated with chronic inflammation of the digestive tract, and primarily includes ulcerative colitis (UC) and Crohn’s disease (CD). The etiology of IBD is not well established but it is usually characterized by inflammation, loss of appetite and weight, chronic diarrhea, bloody stools, fever, rectal bleeding, abdominal pain, fatigue, and anemia (1-3). IBD has been linked to many extra-intestinal manifestations (4) and implicated with mental health problems (5). Current treatments for IBD include, 5-aminosalicylates, glucocorticosteroids, immunomodulators and biologics, and proctocolectomy as a last resort when drug treatment fails. Many of these drugs are often associated with side-effects and various postoperative complications (6, 7). Frontline biologics such as treatment with anti-TNF monoclonal antibodies are only efficacious in some patients and treatment does not result in long term cure (8). Failure of these frontline treatments is associated with elevated risk of colon cancer, and can result in the need for surgical removal of the colon (partial or full). There is therefore an urgent need for new and effective anti-inflammatory drugs to treat IBD.

Guided by millennia of host-parasite co-evolution, we (9-13) and others (14-16) have demonstrated the therapeutic properties of experimental hookworm infection to treat gastrointestinal (GI) inflammatory diseases. Hookworms resident in the human GI tract induce tolerogenic dendritic cells and regulatory T cells which produce suppressor cytokines that keep inflammatory T cells and their effector molecules in check (17, 18). While iatrogenic hookworm therapy shows promise for treating numerous inflammatory diseases in humans, it presents many challenges including apprehension by the patient to readily accept such a radical intervention, safety concerns and regulatory hurdles. In order to circumvent these limitations, we have investigated whether the immunomodulatory properties of hookworms are due to specific metabolites in the parasite’s somatic tissue or excretory/secretory products (ESP). Administration of hookworm ESP to mice protected against inflammation and weight loss in two different mouse models of chemically-induced colitis - the T cell-dependent trinitrobenzene sulfonic acid (TNBS) model (19) and the T cell-independent dextran sulfate sodium (DSS) model (20). We previously characterized the protein constituents (>10 kDa) of hookworm ESP (21), and recently identified a single protein, *Ac*-AIP-2, which in recombinant form displays immunomodulatory properties in a mouse model of asthma that was dependent on regulatory T cells and tolerogenic dendritic cells (DC) (22). A related protein termed *Ac*-AIP-1 was recently shown to protect against inducible colitis by inducing accumulation of regulatory T cells in the mucosa and production of suppressor cytokines including IL-10 and TGF-β (23). Despite progress on the immunoregulatory properties of hookworm ES proteins, much less is known about the composition and anti-inflammatory properties of non-protein small metabolites or low molecular weight metabolites (LMWM; <10 kDa) in hookworm ESP (24).

Other nematodes, namely the parasitic *Ascaris lumbricoides* and free-living *Caenorhabditis elegans*, produce many biologically active LMWM including ascarosides and short chain fatty acids (SCFA) that have diverse biological properties (25). We therefore hypothesized that parasitic nematodes such as hookworms produce LMWM, some of which might have immunoregulatory properties and therefore present as potential anti-inflammatory drug candidates. Using the TNBS model of colitis, we demonstrate that select crude hookworm LMWM extracts afford significant protection against inducible acute colitis in mice and suppress *ex vivo* inflammatory cytokine production from human peripheral blood mononuclear cells (PBMC). Furthermore, we have undertaken both gas chromatography mass spectrometry (GC-MS)- and liquid chromatography mass spectrometry (LC-MS)-guided metabolomics analyses of *A. caninum* total somatic tissue extract and the anti-colitic fractions, and mined these datasets for likely anti-inflammatory candidates.

## RESULTS

To understand the anti-inflammatory role of somatic tissue extracts of adult *A. caninum* and to identify small molecules present in the bioactive extracts, we collected hookworms from dogs, ground them into powder, extracted the LMWM with solvents, and then identified the LMWM present in these extracts using metabolomics techniques. The solvent extractions and preparation of crude extracts were conducted using natural products extraction protocols by sequentially extracting the ground powder with different solvents including mixed solvents of hexane: dichloromethane: acetronitrile (1:1:1 v/v; HDA), followed by dichloromethane (DCM), methanol (MeOH), acidified aqueous (Acidic) and basified aqueous (Basic) solvents. This extraction yielded five somatic extracts (SE) including SE-HDA, SE-DCM, SE-MeOH, SE-Acidic and SE-Basic, which were tested for their anti-inflammatory activities in a mouse model of TNBS-induced colitis and mitogen-stimulated human PBMC.

### Hookworm SE-HDA and SE-MeOH protect mice against clinical symptoms of colitis

Five hookworm somatic tissue extracts were tested for their anti-inflammatory activities using the modified experimental design and TNBS induction in BALB/c mice as previously described by us (26). Extracts (50 μg/mouse) were administered intraperitoneally (i.p.) prior to intra-rectal (i.r.) injection of TNBS and the mice were monitored daily for three days for progression of clinical symptoms of colitis. Of five extracts tested, SE-HDA and SE-MeOH significantly protected mice against TNBS-induced clinical symptoms of colitis including body weight loss, lethargy (mobility), piloerection, diarrhea (fecal consistency) and fecal pellet counts (Fig.1).

**FIG 1.**
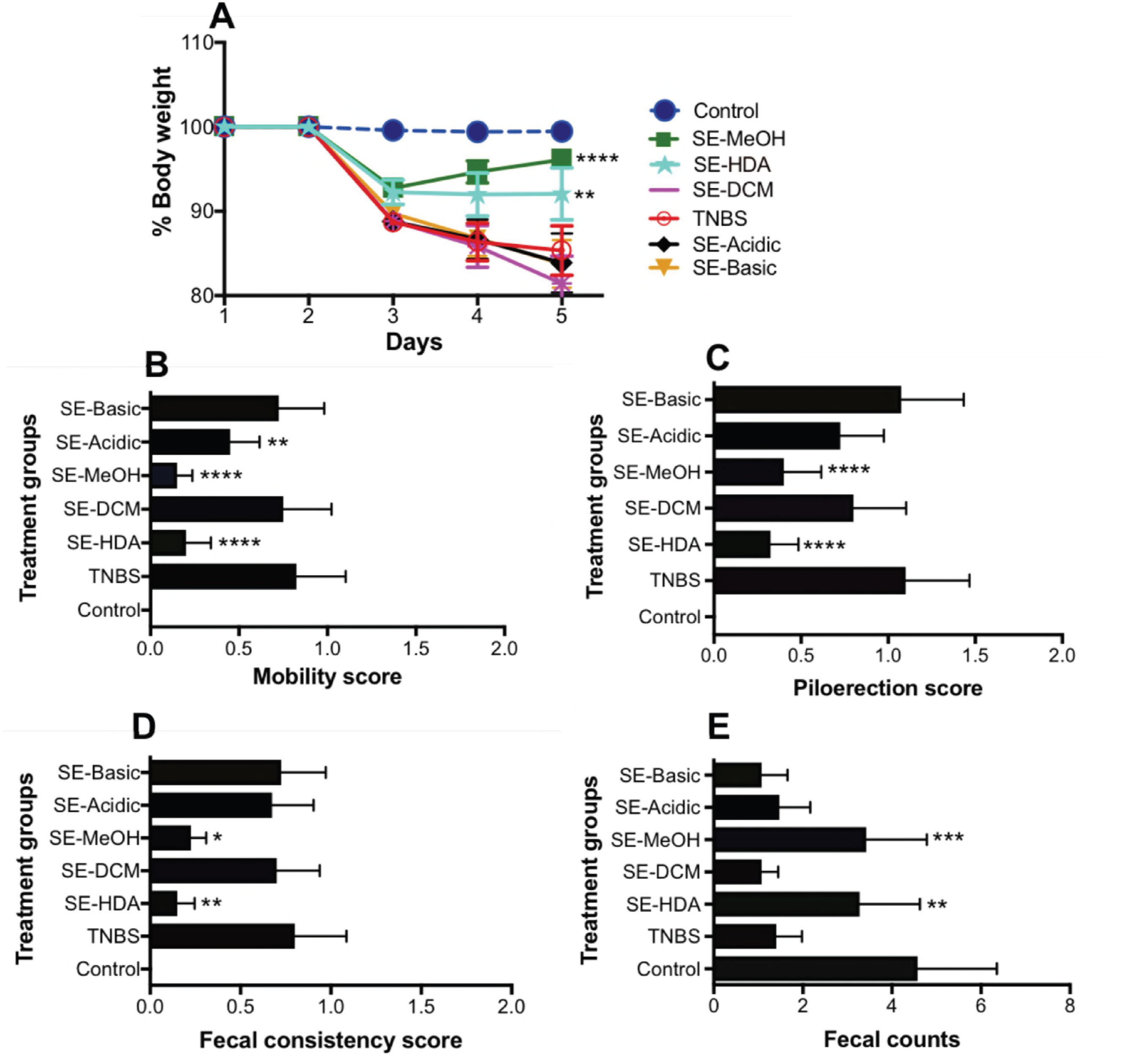
Protective effects of intra-peritoneal administration of somatic tissue metabolite extracts of *A. caninum* against different clinical symptoms of inducible colitis in mice (N = 10). (A) TNBS-induced body weight loss. (B) mobility score. (C) piloerection score. (D) fecal consistency score. (E) fecal pellet counts. Statistical analyses were performed using Graphpad Prism 7 (2way ANOVA and unpaired and nonparametric Mann-Whitney t-test, *P < 0.05, **P *<* 0.01, ***P *<* 0.001, ****P < 0.0001).

### Mice treated with hookworm SE-HDA and SE-MeOH extracts showed reduced pathology

Mice were euthanized on the fifth day of the experiment and scored for pathological progression of colonic inflammation, and colon length was measured. In congruence with the clinical scores, colons of mice treated with SE-HDA and SE-MeOH showed significantly less colonic pathology than those of untreated mice, with significantly longer colon lengths (Fig. 2A), significantly reduced colon thickening (Fig. 2B), fewer adhesions (Fig. 2C), less edema (Fig. 2D), and less ulceration (Fig. 2E).

**FIG 2.**
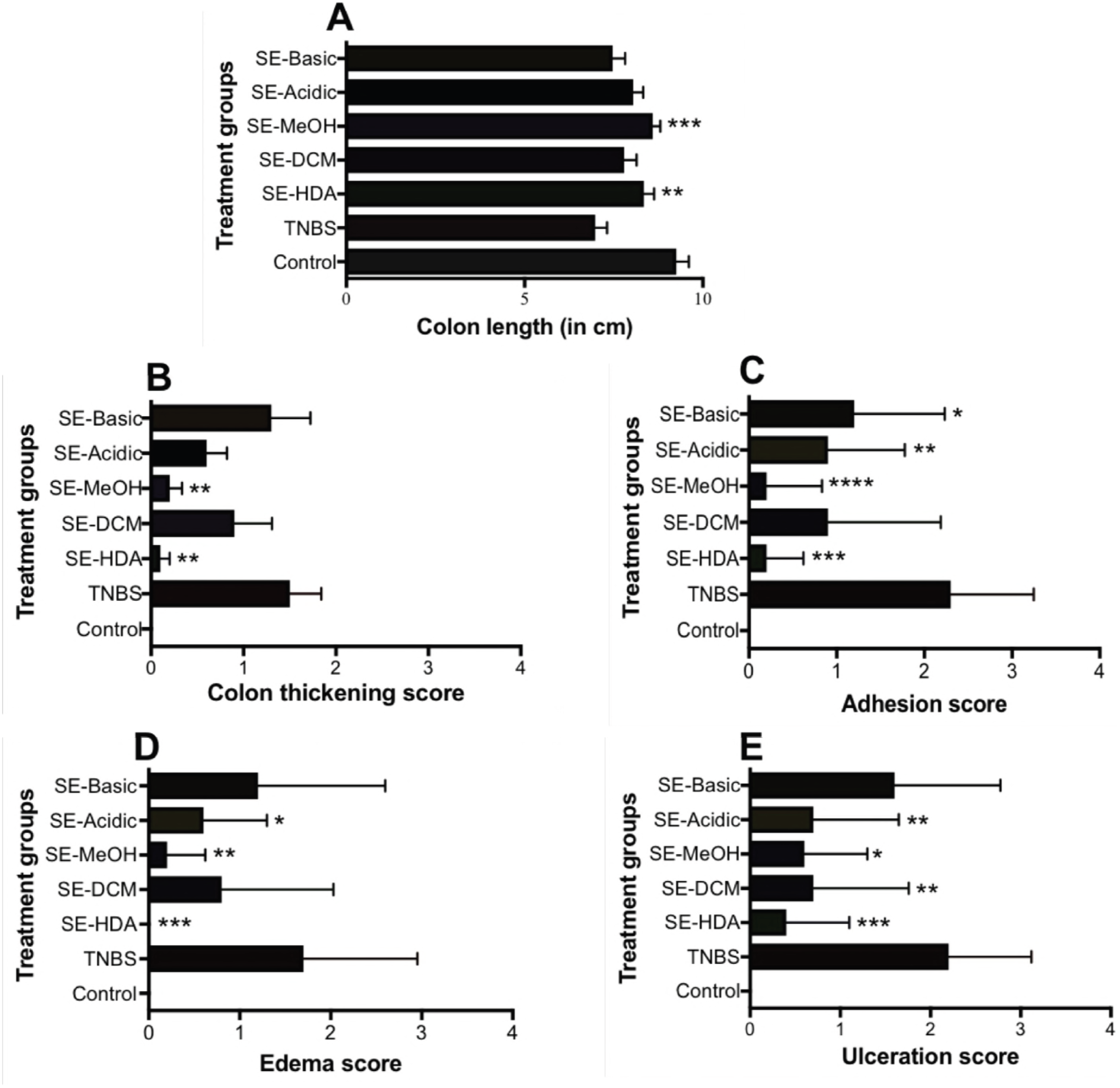
Macroscopic pathology scores of mice treated with different somatic tissue LMWM extracts of *A. caninum* (N = 10). (A) colon length shortening; (B) colon wall thickening; (C) number of adhesions; (D) extent of edema (E); degree of ulceration. Statistical analyses was performed using Graphpad Prism 7 (unpaired and nonparametric Mann-Whitney t-test, *P<0.05; **P<0.01; ***P<0.001; ****P<0.0001).

### Mice treated with hookworm SE-HDA and SE-MeOH showed well-preserved colon architecture

Of five hookworm extracts tested, SE-HDA and SE-MeOH showed well-preserved colon architecture in comparison to the TNBS control group (Fig.3). Naive (healthy control group) mice showed normal colon tissue architecture with healthy crypts and goblet cells, and normal lamina propria and mucosal integrity. Mice that were administered TNBS only (not treated with extracts) developed colitis and exhibited severe thickening of the lamina propria and colon wall musculature, edema, mucosal erosion and destruction of goblet cells. Increased numbers of leukocytes and polymorphonuclear cell infiltrates were clearly evident in the lamina propria and intraepithelial compartments of colons from untreated mice administered TNBS. Treatment with SE-HDA and SE-MeOH prior to administration of TNBS significantly protected against TNBS-induced histopathological damage. Treated mice had well-formed crypts, large numbers of goblet cells, and displayed generally healthy mucosal integrity that was comparable to naive mice that did not received TNBS. The other three hookworm LMWM extracts including SE-DCM, SE-Acidic and SE-Basic did not protect mice against histopathological damage induced by TNBS. Scoring of histological sections for overall colon pathology showed that mice treated with SE-HDA extract had significantly reduced histopathology whereas the SE-MeOH-treated group demonstrated a non-significant trend towards reduced histopathology when compared with TNBS control mice.

**FIG 3.**
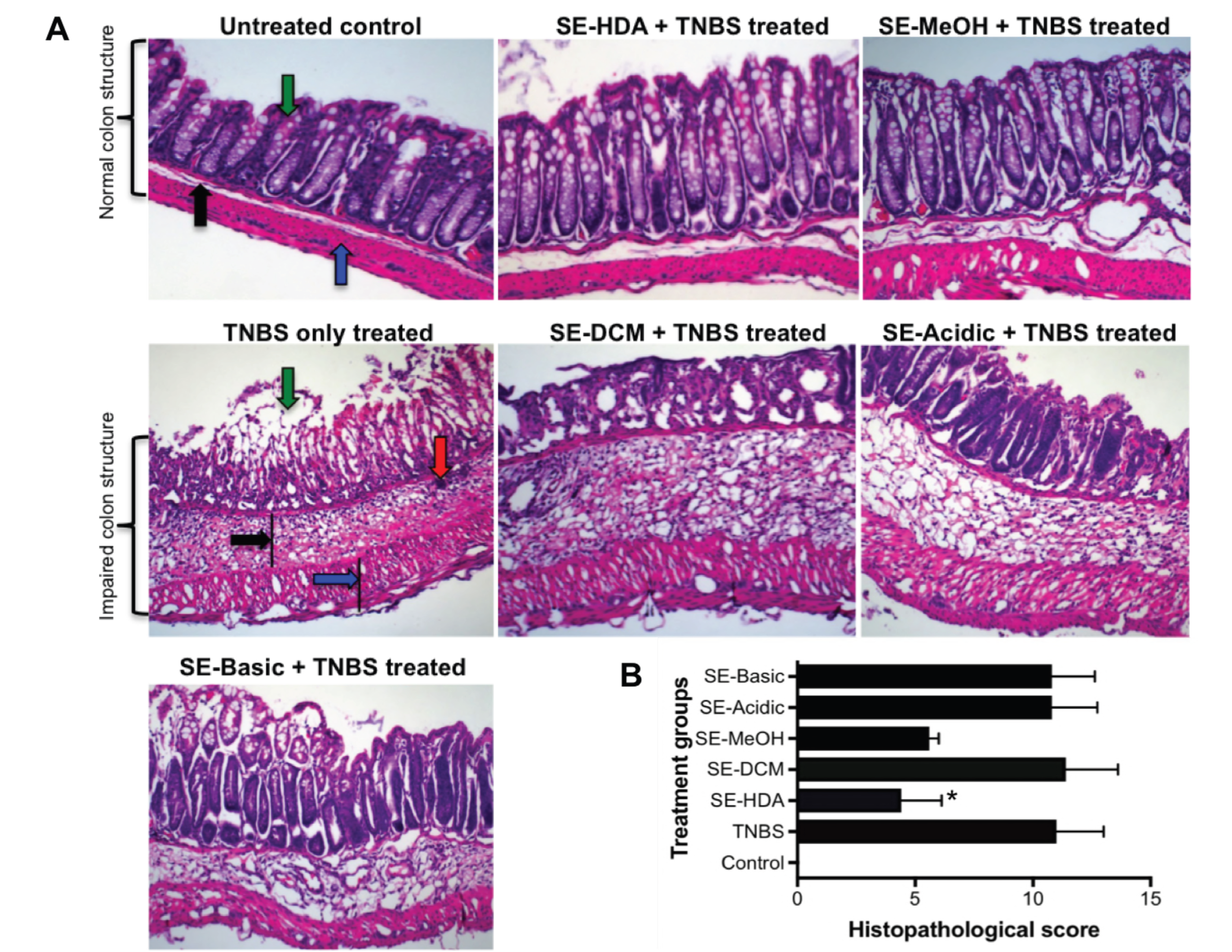
Administration of SE-HDA and SE-MeOH protects mice against TNBS-induced colonic inflammation. (A) Representative histological photomicrographs of haematoxylin and eosin-stained (H/E) paraffin sections (×200) of distal colon tissues of an untreated mouse with normal goblet cells (green arrow), lamina propria (black arrow) and colon wall (blue arrow); TNBS control mouse colon showed erosion of goblets cells (green arrow), thickening of lamina propria (black arrow), cell infiltration (red arrow) and colon wall thickening (blue arrow); mice treated with SE-HDA/TNBS and SE-MeOH/TNBS showed less pathology than the untreated TNBS only control group. Mice treated with SE-DCM/TNBS, SE-Acidic/TNBS and SE-Basic/TNBS extract were not protected against colitic immunopathology. (B) Scoring of histological outcomes of all treatment groups for pathological changes. Statistical analyses were performed using Graphpad Prism 7 (unpaired and nonparametric Mann-Whitney t-test, *P < 0.05).

### Mice treated with hookworm SE-HDA and SE-MeOH extracts had non-significant reductions in colonic inflammatory cytokine production

Mouse colons were surgically removed, cleaned of feces, sliced into 1 cm long pieces, cultured overnight and the supernatants were assessed for colonic cytokine secretion levels. The protection of mice against clinical and pathological changes was consistent with colonic cytokine responses, which showed a non-significant trend towards reduced IFN-γ (P = 0.3538) and IL-17A (P = 0.1510) levels in mice treated with SE-HDA and SE-MeOH (Fig.4).

**FIG 4.**
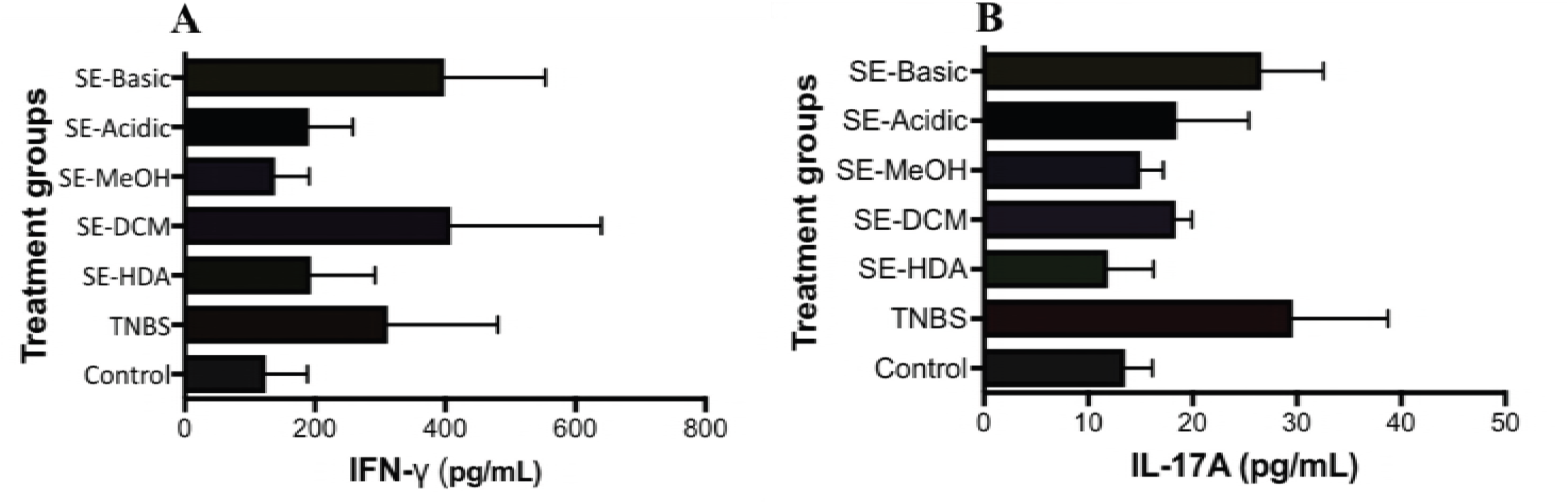
Cytokine profile of mice treated with somatic tissue extracts of *A. caninum* (n = 10). (A) IFN-γ (P = 0.3538). (B) IL-17A (P = 0.1510). Statistical analyses were performed using Graphpad Prism 7 (unpaired and nonparametric Mann-Whitney t-test, P < 0.05 was considered significant).

### SE-HDA suppressed *ex vivo* inflammatory cytokine production by human peripheral blood mononuclear cells

SE-HDA, SE-DCM and SE-MeOH were assessed for anti-inflammatory properties on human PBMC stimulated *ex vivo* with lipopolysaccharide (LPS) to promote cytokine production by myeloid cells or PMA/ionomycin to promote cytokine production by T cells. T cell cytokine production was moderately reduced by SE-HDA at 20 μg/ml including a 14% reduction in IL-2 (P < 0.001), a 34% reduction in IL-6 (P < 0.05), a 59% reduction in monocyte chemoattactant protein-1 (MCP-1) (P < 0.001) and a 33% reduction in TNF-α (P < 0.05) (data not shown) compared to stimulated cells that were not co-cultured with hookworm extracts. We observed a very potent effect of SE-HDA on myeloid cells. LPS-induced cytokines were reduced in the presence of SE-HDA (at 20 μg/ml) including an 88% reduction in IL-1β (P < 0.0001), a 37% reduction in IL-6 (P < 0.01), a 58% reduction in TNF-α (P < 0.01) and an 84% reduction in MCP-1 (P < 0.0001) (Fig. 5).

**FIG 5.**
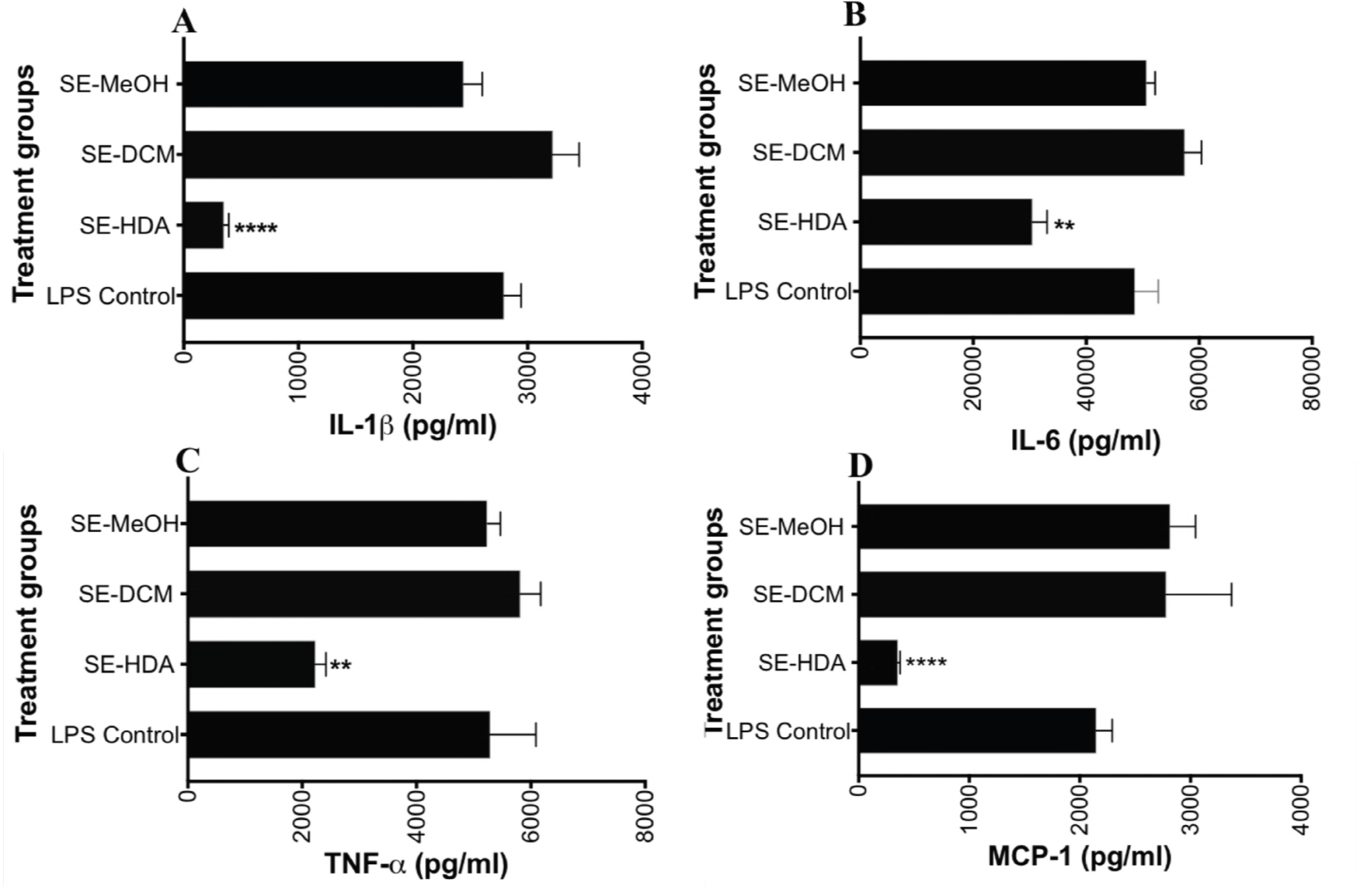
Co-culture of human PBMC with *A. caninum*–derived SE-MeOH, SE-DCM and SE-HDA at a final concentration of 20 μg/ml prior to stimulation with LPS resulted in significant suppression of the pro-inflammatory cytokines IL-1β (A), IL-6 (B), TNF-α (C) and MCP-1 (D). Statistical analyses were performed using Graphpad Prism 7 (unpaired t-test, **P < 0.001, ****P < 0.0001)

A dose-response analysis of the SE-HDA extract at final concentrations of 2, 10, 20 and 50 μg/ml showed that this extract significantly reduced the production of TNF-α, IL-1β, IL-6 and MCP-1 production by LPS-activated PBMC in a dose dependent manner (Fig. 6). The greatest suppression of cytokine secretion was observed for MCP-1, with SE-HDA concentrations as low as 2 μg/ml yielding significantly reduced chemokine secretion. At 50 μg/ml SE-HDA concentration, LPS-stimulated MCP-1 levels were the same as those of unstimulated PBMCs. Analysis of six genetically unrelated individuals showed that SE-HDA suppression of LPS-activated PBMC was consistent with significant reductions in IL-1β, IL-6, IL-12 and MCP-1. A paired analysis of supression in IL-1β and MCP-1 is shown in Fig. 6.

**FIG 6.**
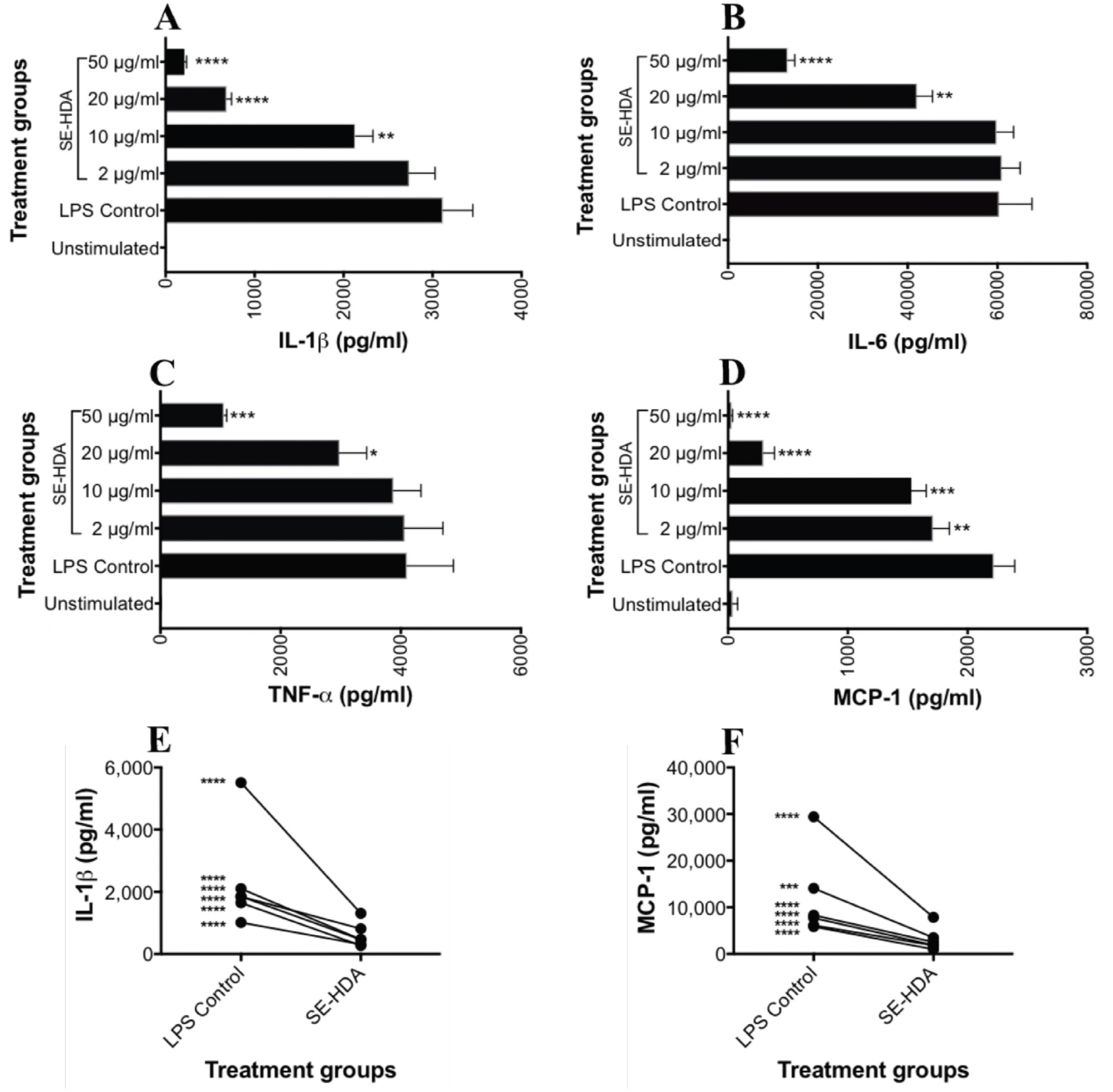
Dose-response effect of inflammatory cytokine and chemokine secretion by human PBMC in the presence of different concentrations of SE-HDA extract of *A. caninum*. Stimulation with LPS resulted in significant suppression of the pro-inflammatory cytokines IL-1β (A), IL-6 (B), TNF-α (C), and MCP-1 (D). A multi-donor analysis showed SE-HDA induced significant suppression of IL-1β (E) and MCP-1 (F) in all 6 donors. Statistical analyses were performed using Graphpad Prism 7 (unpaired t-test, *P<0.05; **P<0.01; ***P<0.001; ****P<0.0001).

### SE-HDA and SE-MeOH somatic tissue extracts contain anti-inflammatory small metabolites

To characterize the small metabolites present in the protective somatic tissue extracts of *A. caninum*, we conducted targeted GC-MS and LC-MS metabolomics analyses of the underivatized SE-MeOH and SE-HDA extracts using methods described previously (27). The compounds were identified by comparing their ion patterns with the ion spectra of the known compounds indexed in the NIST database (28). Through GC-MS analyses, we identified a total of 32 metabolites from these two bioactive somatic tissue extracts (Table S1). Each extract contained 20 metabolites, with 12 of them found exclusively in each extract. Both extracts contained eight medium to long chain free saturated and unsaturated fatty acids (C_11_ to C_26_). Palmitic acid, methyl palmitate, stearic acid and methyl stearate were abundant in both extracts. The most common mono/poly-unsaturated fatty acids present in both crude extracts were oleic acid, linoleic acid, γ-linolenic acid and palmitoleic acid. Six of these fatty acids including stearic acid (C18:0) (29), palmitic acid (C16:0) (30), methyl palmitate (31), γ-linolenic acid (32), palmitoleic acid (C16:1n-7) (33) and oleic acid (C18:1) (29, 34) have been reported to exhibit anti-inflammatory properties.

Using LC-MS, we identified eight SCFA, with isovalarate as the major component present in both the SE-MeOH and SE-HDA somatic extracts. Acetate, propionate, butyrate, 2-methylbutyrate, isovalerate, caproate and heptanoic acid were present in both protective fractions/extracts (Table S2). Isobutyrate was present only in the SE-MeOH extract. Acetate, propionate and butyrate (obtained from other synthetic sources) were previously reported to have anti-inflammatory activities (35).

### Global metabolomes of derivatized whole worm somatic tissue extract of hookworms

Metabolomics analyses of two protective extracts (SE-HDA and SE-MeOH) showed only the representative metabolomes of underivatized samples of hookworm extracts. To gain a global insight of small metabolites present in *A. caninum*, we conducted a targeted GC-MS and LC-MS metabolomics analyses of the methoximated, trimethylsilyl derivatized whole worm extract. Whole worm extracts were divided into polar and non-polar fractions. From the polar fraction, using GC-MS we identified 47 polar small metabolites (Table S3) belonging to seven chemotypes including amino acids, sugars, sugar phosphates, organic acids, glycerides, carbamides and oligosaccharides. Glycine, L-valine, L-proline, pyroglutamic acid, L-isoleucine, L-tryptophan, D-talose, D-glucose, L-lysine and γ-aminobutyric acid (GABA) were present in abundance. Eleven of these polar metabolites including glycine (36), L-isoleucine (37), L-lysine (38), γ-aminobutyric acid (GABA) (39), mannitol (40), D-ribose (41), trehalose (42), L-histidine (43), uridine (44), L-methionine (45) and citric acid (46) have been previously shown to exhibit anti-inflammatory activities against various diseases, including arthritis, renal inflammation, subarachnoid hemorrhage, pulmonary fibrosis and other conditions.

From the non-polar fraction, using GC-MS we identified 22 fatty acids (Table S4) comprising 17 saturated fatty acids (including stearic acid, palmitic acid, arachidic acid, margaric acid and myristic acid) and five unsaturated fatty acids (including elaidic acid, oleic acid, erucic acid, petroselinic acid, and nervonic acid). Literature review and content analyses revealed that at least seven of these fatty acids including stearic acid (C18:0) (29), palmitic acid (C16:0) (30), methyl palmitate (31), lauric acid (C12:0) (47), capric acid (C10:0) (47), γ-linolenic acid (32) and caprylic acid (C8:0) (48) were previously reported to have anti-inflammatory activities.

Using LC-MS analysis, we identified five SCFA from whole worm somatic tissue extracts, including isobutyrate, propionate, 2-methylvalerate, acetate and butyrate (Table S2). Of these SCFA, acetate (C2:0), propionate and butyrate (C4:0) have been reported to be effective in preventing inflammation associated with IBD (35).

## DISCUSSION

Parasitic helminths, such as hookworms, have evolved to establish chronic infections in the human gut while inducing minimal pathology when present in low numbers (17). Hookworms regulate the immune system of the host for their benefit and subsequently promote a state of immune tolerance. This regulated environment not only promotes longevity for the parasites but also reduces the likelihood of the host developing diseases that result from a dysregulated immune system (49). A significant number of experimental and clinical studies support the immunoregulatory proficiency of parasitic helminths against distinct immunopathologies, including allergies, IBD and other autoimmune diseases (9, 16, 50-53). While iatrogenic infection with hookworms and other helminths shows promise for treating numerous inflammatory diseases in humans, the therapy presents many challenges including patient apprehension, safety concerns and regulatory hurdles (17, 54). We and others showed that *A. caninum* ES proteins (>10 kDa) have potent immunomodulatory properties and can protect mice against pathology in different inducible models of colitis (17, 19, 20, 55, 56) and asthma (22). Non-proteinaceous small metabolites derived from helminths, however, have received far less attention in terms of their molecular characterization and their immunoregulatory properties. Indeed, we recently proposed that helminth LMWM warrant in-depth investigation as an untapped source of new drugs for treating inflammation (24). Here we provide a molecular characterization of hookworm LMWM and assess the efficacy of five different LMWM extracts in preventing both the onset of inducible colitis in mice and inflammatory cytokine production by primary human leukocytes.

Considering the relative strengths and limitations of all the available animal models, we chose the TNBS colitis model, which represents an intestinal immune response to environmental triggers, to investigate the anti-inflammatory properties of hookworm somatic LMWM. The TNBS model induces a mixed Th1/Th2/Th17 response (8), facilitating its use in screening for drugs targeting a broad range of inflammatory pathways. The intestinal mucosa of a TNBS-treated mouse is characterized by rapid production of inflammatory cytokines such as IFNγ and IL-17A (52). These cytokines are particularly important because disruption of the colonic mucosal layer by ethanolated TNBS dysregulates intestinal goblet cell function and elicits inflammation that drives the production of these cytokines. The colon contains epithelial goblet cells, which are instrumental in controlling intestinal immune homeostasis by producing a protective mucus layer. Therefore, any extracts/small molecules from intestinal worms that can promote retention of colonic mucus and inhibit inflammatory cytokine production have therapeutic potential. We showed that mice treated with *A. caninum* LMWM extracts (SE-MeOH and SE-HDA) promoted retention of healthy gastrointestinal architecture after TNBS administration; this took the form of normal mucosal crypts, large numbers of unaltered goblet cells, and normal lamina propria and colon wall architecture in comparison to untreated groups that received TNBS. The colon culture supernatants and homogenates of these two treatment groups showed trends towards reduced expression (albeit non-significant) of TNBS-induced inflammatory cytokines, including IFN-γ and IL-17A. The protection conferred by SE-MeOH and SE-HDA extracts against TNBS-induced colitis was, however, more evident in their ability to significantly reduce body weight loss, improve clinical IBD related symptoms (including abridged mobility, piloerection and fecal consistency), and arrest pathological progression (including colon length shortening, colon wall thickening, adhesion, edema and ulceration). However, when these two anti-colitic somatic tissue extracts were assessed *ex vivo* for suppression of LPS-activated human PBMC, only treatment with SE-HDA resulted in significant suppression of pro-inflammatory cytokines including TNF-α, IL-1β, IL-6 and the chemokine MCP-1. Despite conferring robust protection against TNBS-induced colitis in mice, SE-MeOH did not suppress the production of LPS-stimulated cytokines by human PBMC. This could be due to distinct mechanisms of action of the bioactive components in each of the two models, or might also reflect the presence of different bioactive metabolites in the two distinct fractions. SE-HDA-induced suppression of inflammatory cytokines and chemokines was dose dependent, and particularly potent for MCP-1. MCP-1 and other chemotactic cytokines such as IL-8 induce chemotaxis, leukocyte activation and granule exocytosis, all of which increase chronic inflammation and intestinal tissue destruction in IBD (57).

The GC-MS and LC-MS analyses of these two anti-colitic somatic extracts (SE-MeOH and SE-HDA) revealed the presence of 32 small metabolites including eight medium-long chain fatty acids and seven SCFA. Some SM detected here such as ficusin, naphthalene derivative, methyl behenate and hentriacontanes are compounds known only from plants, and they are unlikely to be synthesized by nematodes. Their putative appearance in these two helminths suggest the dietary exposure of the host animals from which the hookworms were obtained. These SM are tentative assignments pending MS-MS/NMR confirmation of proposed structures. From the whole worm somatic tissue extract of *A. caninum*, we identified 47 polar metabolites (glycine as the major amino acid), 22 fatty acids (stearic acid and methyl palmitate as major components) and four SCFA (isovalerate as the major SCFA). The presence of a large number of polar metabolites was consistent with the known dependence of these parasites on glucose metabolism, glycogen synthesis, oxidative metabolism and phosphorylation (58-62). Palmitic acid and stearic acid were the most common saturated fatty acids present in all the hookworm extracts. Palmitic acid is the primary fatty acid from which other longer fatty acids are synthesized. Linoleic acid is found mostly in plant glycosides and is used by animals in the biosynthesis of prostaglandins (via arachidonic acid) and cell membranes (63). SCFA, including acetic, propionic, methylbutyric, *n*-valeric and methylvaleric acids, were first reported in 1965 (64) from the cuticle, muscle and reproductive systems of *A. lumbricoides*. Our findings concur with previous findings (62) on the presence of SCFA in the ES products of *A. caninum*, where volatile SCFA such as acetic, propionic, isobutyric and branched chain C_5_ acids were detected (62). More recently, saturated fatty acids were reported from the ova of *A. caninum* (65), but the authors did not report the presence of SCFA.

The nature and origin of these excreted SCFA of *A. caninum* were demonstrated through carbon (D-glucose-^14^C) isotope/radiocarbon labeling, and indicated the presence of a glucose metabolism intermediate between that observed in aerobes and that characteristic of helminth anaerobes (62). The importance of SCFA as intermediates in the energy metabolism of intestinal parasites in a succinate decarboxylase-dependent manner, as well as uncoupled mitochondrial respiration, has been reported (66). A recent review (67) suggested that formation and excretion of acetate as a metabolic end product of energy metabolism occurs in many protist (*Giardia lamblia*, *Entamoeba histolytica*, *Trichomonas vaginalis, Trypanosoma* and *Leishmania* spp.) and helminth (*Fasciola hepatica*, *Haemonchus contortus* and *Ascaris suum*) parasites from acetyl-CoA by two different reactions, both involving substrate level phosphorylation that is catalyzed by either a cytosolic acetyl-CoA synthetase or an organellar acetate:succinate CoA-transferase. However, these enzymes involved in SCFA biosynthesis were poorly represented when we mapped them against the known metabolic pathways (KEGG) of 81 worm genomes including the human hookworm, *Necator americanus*, and the model free-living nematode, *C. elegans*. None of the hookworm species represented in genome databases have any annotation associated with SCFA synthesis other than two enzymes involved in propanoate synthesis. It is known that parasitic helminths modulate intestinal inflammation *via* alteration of the composition of the gut microbiota. This mechanism has been demonstrated both in rodent models (68) and human studies where iatrogenic hookworm infection resulted in increased bacterial species richness and elevated production of SCFA (11, 69). It is apparent that the gut microbiome is a complex ecosystem with microbial syntrophy at the intestinal mucosal interface (70), and that SCFA such as acetate, butyrate and propionate are produced and utilized by bacteria, and benefit host epithelial cells by producing molecules like vitamin B_12_ (71). SCFA are key metabolites and energy sources for commensal bacteria at the gut interface. Therefore, while it is possible that *A. caninum* synthesizes SCFA *de novo*, further studies will be needed to define the contribution of the commensal microbiome to SCFA synthesis in hookworms.

Analyses of the available literature on the biological activities of all the small molecules identified from the somatic tissue extracts of *A. caninum* showed that 11 polar metabolites (36-46), nine medium-long chain fatty acids (29-34, 47, 48), and three SCFA (35) have been previously isolated from different sources and have been found to exhibit anti-inflammatory activities. Of 23 anti-inflammatory small metabolites, only three compounds including one polar metabolite (glycine) and three SCFA (acetate, propionate and butyrate) were studied *in vivo* against chemically- and stress-induced ulcers in the gastric mucosa (36). Fatty acids, especially SCFA, have roles in host defense against potential opportunistic or pathogenic microorganisms, and establish an immunoregulatory environment that protects against inflammation in both the gastrointestinal tract and distant sites including the lung and heart (68, 72-74). SCFA are particularly important for colon homeostasis (72). For example, butyrate nourishes the colonic mucosa, and butyrate administration confers beneficial effects against IBD (72, 75). A comparative *in vitro* study of acetate (C2:0), propionate and butyrate (C4:0) revealed that propionate and butyrate were equipotent, where as acetate was less effective against IBD (35). Butyrate and propionate have also been implicated in the maintenance of host immune function by signaling to epithelial cells, maintaining regulatory T cell populations and inhibiting macrophage activation (70, 71). It is possible that *A. caninum* may be actively skewing the microbiome towards an anti-inflammatory composition by enhancing tolerogenic immune crosstalk through SCFA production. Chronic infection with *H. polygyrus* altered the intestinal habitat, resulting in increased SCFA production that was transferrable via the microbiota alone, and was sufficient to mediate protection against allergic asthma (68). The resulting anti-inflammatory cytokine secretion and regulatory T cell suppressor activity required the G protein-coupled receptor (GPR)-41, further highlighting the essential role of SCFA produced by commensal microbe communities shaped by the presence of helminth infections.

In summary, this study demonstrates that the somatic LMWM extracts of *A. caninum* are diverse in nature and possess anti-inflammatory properties that can suppress colitis in mice and inflammatory cytokine production by human leukocytes. Of five extracts tested, SE-MeOH and SE-HDA significantly protected mice against chemically-induced colitis, which implied that methanol and HDA were the best solvents to extract bioactive metabolites from hookworm somatic tissues. Our GC-MS and LC-MS analyses highlighted the presence of multiple small molecules in these fractions. Several of these metabolites, including the SCFA, have been previously shown to have anti-inflammatory properties in various target diseases, including IBD. It is possible that these anti-inflammatory small molecules, either individually or in synergy, are responsible for the anti-inflammatory properties of SE-MeOH and SE-HDA extracts of *A. caninum.* Future work will entail purification and isolation of the bioactive metabolites, synthesis of the candidate components, and detailed pharmacotherapeutic assessment of the anti-colitic properties of the candidate compounds using a chronic immunologic mouse model of colitis. Moreover, while it is difficult to obtain large quantities of hookworm ESP, efforts should be invested in further isolating the ESP metabolomes with a focus on understanding the source and nature of SCFA, given that these compounds have specifically evolved to interact with host tissues and play important roles in regulating inflammation.

## MATERIALS AND METHODS

### Hookworm collection

*A. caninum* adult worms were collected with clockmaker tweezers from the small intestine of infected dogs and transferred to pre-warmed culture media (2% Glutamax in phosphate buffered saline (PBS), 5% antibiotic/antimycotic [AA]) in 50 ml falcon tubes. *A. caninum* adult worms were thoroughly washed with 5× antimycotic/antibiotic solution to purge bacterial contaminants and were cultured (50 worms per dish) in a single component Glutamax medium (2% Glutamax in PBS supplemented with 2× AA) for 2 h at 37°C in 5% CO_2_ to allow regurgitation/defecation of the host blood meal and other material (including bacteria) that may have been acquired from the dog host gut. The worms were then snap-frozen in liquid nitrogen until further use.

### Preparation of hookworm somatic extracts

The adult worms were frozen with liquid nitrogen and made into powder using a mortar and pestle. The powder was first soaked in a solvent mixture of hexane:dichloromethane:acetonitrile (1:1:1 v/v, 6 ml/g; HDA) for 30 min and was filtered (Whatman 4, Qualitative circles 185 mm, England International Ltd.). The extraction of cell material on filter paper was repeated three times with the same solvent. The filtrates were combined, centrifuged at 1,831 *g* (Rotina 420 R, Hettich Zentrifugen, Germany) for 20 minutes at 4°C and the supernatant slowly transferred to round-bottom flasks. The solvent was removed using a Rotary Evaporator (G5 Heidolf CVC 3000 Vacuubrand) to obtain the somatic tissue HDA fraction (SE-HDA). The solid residue was snap-frozen in liquid nitrogen, powdered and extracted with dichloromethane (3×). The filtration, centrifugation and drying processes were repeated as above to obtain the DCM fraction (SE-DCM). The remaining somatic tissue was snap-frozen again, powdered and extracted with methanol to obtain the MeOH fraction (SE-MeOH). Remaining solid residue from this step was snap frozen and soaked in 5% HCL (pH 1-2). The supernatant was freeze-dried using a Scanvac Cool Safe to obtain acidified aqueous extract (SE-Acidic). The solid tissues were finally soaked in basified (NH_4_OH, pH 10-12) water, filtered and centrifuged at 1,831 *g*, and freeze-dried to obtain alkaline aqueous extract (SE-Basic). Stock concentrations of these extracts (1 mg/ml) were prepared by dissolving 1 mg of each extract in 20 μL of DMSO followed by addition of 980 μl of PBS. From each stock concentration, 50 μl was transferred to a vial and added to 150 μl of fresh PBS to make a total injectable volume of 200 μl/mouse. These extracts were assessed for efficacy in the TNBS model of acute colitis.

### Animal ethics, source and housing of mice

The James Cook University (JCU) animal ethics committee approved all experimental work involving animals (Ethics approval number A2199). Mice were raised in cages in the JCU animal facility in compliance with the Australian Code of Practice for the Care and Use of Animals for Scientific Purposes, 7^th^ edition, 2007 and the Queensland Animal Care and Protection Act 2001. Age-matched 5-week old male BALB/c mice were sourced from Animal Resources Centre (Perth, Australia), weighed, placed in cages (5 mice per cage), and allowed to settle into the study facility for 4-5 days prior to the start of the experiment. Animals were housed in a temperature (26°C) and humidity-controlled (40-70%) environment, exposed to a 12-hour day/night cycle, and provided with irradiated mouse chow supplied by Specialty Feeds (Glen Forrest, Western Australia) and autoclaved tap water *ad libitum*.

### Experimental design and induction of TNBS colitis

We followed the modified experimental design and TNBS induction protocols described by us earlier (26). Mice were divided into three different groups as “Naïve”, “TNBS only” and “TNBS + Sample treatment”. This experiment was performed in duplicate (total n=10 mice/group). The samples were filtered using 0.22 μM sterile filters prior to intra-peritoneal (i.p) administration to mice. Each mouse in the “Sample treatment” groups (each group labelled as SE-HDA, SE-DCM, SE-MeOH, SE-Acidic and SE-Basic) received 50 μl of extract suspension/mouse by i.p. injection (Day 1). The TNBS only group was administered 200 μl of PBS/DMSO (1%). Twenty-four hours post sample administration (Day 2), the mice were anaesthetized by i.p. injection of anesthetic solution containing 50 mg/kg of ketamine and 5 mg/kg of xylazine. TNBS was administered by intra-rectal (i.r.) injection of 100 μl TNBS (2.5 mg/mouse) mixed with 50% ethanol using a soft catheter (Insyte Autoguard Shielded IV catheter 20G × 1” Pink; Becton Dickinson). Mice were kept inverted for 2 minutes to prevent leakage of TNBS and returned to their cages. Mice were monitored daily until euthanized at day 5.

### Monitoring colitis progression by clinical scoring

Mice were clinically scored for weight loss, piloerection, mobility, fecal consistency and fecal pellet counts for 3 days following administration of TNBS. Each individual mouse was placed in a clean cage/open plastic jar and observed for 10 minutes to score the clinical symptoms. The changes in clinical signs of disease were scored from 0-2, 0 being normal and 2 being diseased. A score of “0” was given when the mice gained weight, “1” when weight remained the same and “2” upon losing weight. Mice with no piloerection scored “0”, mild piloerection over the neck as “1” and severe piloerection all over the body as “2”. The mobility of a mouse was scored as “0” for normal, “1” for movement only after stimulation, and “2” for hunched posture with no movement around the cage, even after stimulation. For fecal consistency, normal feces were scored “0”, mild diarrhea was scored “1” and severe diarrhea with blood or no feces was scored “2”. After 10 minutes of isolation, the fecal pellets were counted (higher number of pellets indicated normal or mice recovering from chemical colitis).

### Assessing intestinal pathology and macroscopic scoring

On day 5 post-TNBS administration, the mice were euthanized. Their colons (from cecum to rectum) were surgically removed, measured and assessed for changes in macroscopic appearance and pathological parameters including adhesions, bowel wall thickening and edema (scoring matrices of 0–3; 0 = normal, 1 = mild, 2 = moderate, and 3 = severe). The colons from each group were lined up on a clean surgical drape paper towel, their lengths measured and then transferred to a petri dish in sterile DPBS. The tissues were opened longitudinally, washed with DPBS, placed under a microscope (Olympus SZ61, 0.67–4.5×), observed for inflamed sections, and scored for ulceration (0 – 3).

### Evaluating colon histological structure

The distal colon tissue sections (1 cm) that were harvested from mice were placed in 4% paraformaldehyde (1?ml) to fix tissue overnight at 4°C and then transferred to 70% ethanol for storage. Tissue was embedded in paraffin and sectioned longitudinally for histology at 4 μm thickness. Sections were stained with hematoxylin and eosin (H/E), observed for histological changes by light microscopy and histological photomicrographs (×200) were captured. Each histological photomicrograph was blinded and then scored for changes in overall colon morphology and epithelial integrity. The cross-sections of the colon tissues were scored for inflammation, edema, hyperplasia, ulceration and number of goblets cells using a scoring matrix (76). An AxioCam Imager –M1 (MRC ZEISS) was used for scoring the colon histology cross-sections.

### ELISAs and cytokine measurement of colon tissues

Colon pieces (1?cm) were collected and placed in sterile 24 well tissue culture plates with 500?μl of complete medium (RPMI containing 10% fetal bovine serum, 1% penicillin/streptomycin, 0.1% β-mercaptoethanol, 1% HEPES buffer. Tissues were cultured for 24?h at 37°C (supplied with 5% CO_2_) after which supernatants were collected and stored at −80°C until further analysis. Colon culture supernatants were thawed and levels of cytokines were quantified using a sandwich enzyme-linked immunosorbent assay (ELISA) (Ready-SET-Go!, eBiosciences) following the manufacturer’s instructions. OD_490_ values were measured using a POLARstar Omega plate reader (BMG LABTECH) and were expressed as picogram (pg) of tissue weight per mL.

### Human PBMC collection and culture conditions

The human blood used for this study was obtained from healthy volunteer donors. Written informed consent was obtained from each donor at the time of blood draw. Ethical approval for this research was obtained from the James Cook University Human Ethics Committee. PBMC were isolated from whole blood by density gradient centrifugation using Ficoll-Paque media. For induction of T cell cytokines, PBMC were activated with a cell stimulation cocktail of 50 ng/ml of phorbol 12-myristate 13-acetate (PMA) and 1 μg/ml of ionomycin (eBioscience). PMA + ionomycin-stimulated cells were treated with 20 μg/ml of hookworm extracts (SE-HDA, SE-DCM, SE-MeOH) or remained untreated. For stimulation of myeloid-associated cytokines, PBMC were activated with 10 ng/ml lipopolysaccharide (LPS) (Sigma-Aldrich). LPS-stimulated PBMC were treated with 2-50 μg/ml of hookworm extracts (SE-HDA, SE-DCM, SE-MeOH) or remained untreated. The cell culture plates were incubated overnight at 37°C and 6.5% CO_2_. After incubation, the samples were centrifuged at 1,500 *g* for 5 minutes and the culture supernatants were collected for cytokine analysis.

### BD™ Cytometric Bead Array

IL-1β, IL-6, IL-12, TNF-α and MCP-1 from PBMC culture supernatant were quantified using a cytometric bead array (CBA) (BD^™^ Biosciences). The CBA assays were performed according to the manufacturer’s instruction and run using a five laser Special Order LSRFortessa^™^ with HTS (BD Biosciences). Cytokine concentrations (pg/ml) were calculated based on the sample MFI compared to the cytokine standard curves. BD™ FCAP Array software version 3.0 was used for data analysis. Graphs and statistical analysis were produced using GraphPad Prism version 7.02 (GraphPad Software Inc).

### Cryomill somatic tissue extraction of adult worms for GC-MS metabolomics analysis

Somatic tissue extract of adult *A. caninum* (10-20 mg) was snap-frozen in liquid nitrogen to arrest metabolic changes, placed in cryomill tubes, then suspended in 600 μl extraction solution (methanol:water 3:1 (v/v) containing internal standards ^13^C,^15^N-valine and 13C-sorbitol). The sample was extracted using a Precellys 24 Cryolys unit (Bertin Technologies) at 6800 rpm, 3 × 30 sec pulses, 45 second interval between pulses, temperature < −5°C (pre-chilled with liquid nitrogen). The homogenate was transferred to a fresh microfuge tube on ice and chilled chloroform (150 μl) was added. The solution (chloroform:methanol:water 1:3:1 (v/v) monophasic mixture) was vortexed vigorously chilled on ice for 10 minutes with regular mixing and then centrifuged for 5 minutes at 0°C. The supernatant was transferred to a fresh 1.5 ml microfuge tube on ice and milli-Q water (300 μL) was added to each tube to obtain a biphasic partition of the solution (chloroform:methanol:water 1:3:3 (v/v)). The sample was vortexed vigorously and then centrifuged at 0°C for a further 5 minutes.

For total fatty acid analyses, the bottom chloroform extract (45 μl) was transferred to fresh tubes and 0.2M *m*-trifluromethylphenyl trimethyl ammonium hydroxide (methprep) (5 μl) was added. Samples (5 replicates) were mixed using a thermomixer at 750 rpm for 30 minutes at 37°C. The samples were injected (2 μl) into an Agilent 7000 triple quad GC-MS (1:10 split injection, BPX70 60 m × 0.25 mm × 0.25 μm column) and the raw data were obtained and processed.

### Derivatization method for targeted metabolite analysis using GC-MS

The upper aqueous phase (methanol:water, ~900 μl) was transferred to a fresh tube, then aliquoted (50 μl) into a pulled point insert and dried in an evaporator (Christ RVC 2-33 CD, John Morris Scientific Australia) at 35°C. A further 50 μl of the aqueous sample was added every 30-40 minutes until completely dry. Samples were dehydrated by adding 50 μl anhydrous methanol, and then derivatized by addition of 20 μl methoxyamine (30 mg/ml in pyridine, Sigma Aldrich/Merck) at 37°C for 30 minutes, and then 20 μl of *N*,*O*-Bis(trimethylsilyl) trifluoroacetamide (BSTFA) + 1% trimethylchlorosilane (TMCS) (ThermoFisher Scientific) was added prior to incubation at 37°C for 30 minutes. The derivatized sample (1 μl) was analyzed using an Agilent 7890 GC-MS (5973 MSD) (77). Chromatographic separation was achieved using an Agilent VF-5 ms column (30 m × 0.25 mm × 0.25 μm). Conditions for the oven were set at 35°C, held for 2 minutes, then ramped at 25°C/minute to 325°C and held for 5 minutes. Helium was used as the carrier gas at a flow rate of 1 ml/minute, with compounds being detected across the m/z range of 50–600 atomic mass unit (amu).

### GC-MS analyses of SE-HDA and SE-MeOH extracts using underivatized protocols

We performed GC-MS analysis on the two bioactive extracts using methods described by us previously (78). The dried crude fractions were re-suspended in chloroform-methanol solvent (90%:10%) and were directly injected into a Shimadzu GC-2010 Plus system to analyse their chemical constituents. The GC system used helium as a carrier gas (1.22 mL/min, pressure 67.7 kPa at 40°C in a constant total flow mode) and the separation was achieved using a DB-5 ms column (l30 m length × 0.25 mm, i.d., 0.25 μm, Phenomenex). Injector (injection – splitless mode) and detector temperatures were set at 225°C and 300°C, respectively. The starting oven temperature was programmed at 45°C with an increasing temperature of 3°C/minute until it reached 100°C (hold time = 4 minutes) and final temperature of 240°C (hold time = 50 minutes). Similarly, the same equipment was programed for the MS system (condition and column same as above) with a runtime of 90 minutes (ion source = 200°C, solvent cut time = 1 minute, threshold = 0, starting mass (*m*/*z*) = 35 and maximum mass measured (*m*/*z*) = 1000, acquisition mode = scan, scan speed = 3333). The GC-MS was acquired in scan mode producing a total ion chromatogram (TIC).

The chemical constituents were identified by comparing and matching the ion fragmentation patterns of the test sample compounds with the National Institute of Standards and Technology (NIST, USA) mass spectra library of GC-MS data. Each compound was then surveyed for erstwhile literature on their anti-inflammatory properties using SciFinder Scholar, PubMed and Google Scholar. Published studies reporting anti-inflammatory activities for each compound are cited in the relevant tables.

### Identification of SCFA using LC-MS protocols

The SCFA in the protective SE-HDA and SE-MeOH extracts, and the whole worm extract of hookworms were analysed by LC-MS in accordance with previously described protocols (79). Samples were analyzed in triplicate. Known SCFA, including acetate, propionate, isobutyrate, butyrate, 2-methylbutyrate, isovalerate, valerate, 3-NPH, 2-methylvalerate, 3-methylvalerate, isocaproate, caproate and heptanoic acid were used as standards (5 μM and 50 μM concentrations). These SCFA were mapped against the existing 81 worm genome KEGG pathways to understand their biosynthetic nature.

### Statistical and data analyses

The data from groups of mice from 2 independent experiments (N =10) were combined and the statistical analyses were performed using GraphPad Prism (version 7.0) as described earlier (76). Comparisons were made between the sample treatment + TNBS groups and the TNBS only group; P values of < 0.05 were considered significant. When determining the differences between more than two groups with equal numbers of mice/samples, 2-way ANOVA was used. When two groups were compared a Mann-Whitney (unpaired, non-parametric) t-test was applied. All results reported denote mean ± standard error of the mean (SEM). The metabolomics data was analyzed in a targeted approach using Agilent’s Mass Hunter Quantitative Analysis software (v.7). Target ion peak areas for polar and non-polar metabolites were extracted using the in-house Metabolomics Australia metabolite library and were integrated and output as a data matrix for further analysis (section 3.5).

## ACKNOWLEDGMENTS

This study was supported by a NHMRC Peter Doherty Early Career Researcher Fellowship (APP1091011) and AITHM Capacity Development Grant to P.W; a NHMRC program grant APP1037304 and NHMRC Senior Principal Research Fellowship (APP1117504) to A.L; and a NHMRC Career Development Fellowship (APP1131732) to J.J.M.

We thank Dr. Makedonka Mitreva for hookworm genome analyses, Dr. Severine Navarro, Mr. Atik Susianto, Dr. Komal Kanojia, Dr. Vinod Narayana and Mr. Bjoernar Hauge for assistance with cytokine analyses, SCFA method development for hookworm extracts and animal husbandry.

